## Supplement_files for "Brochosomes as an antireflective camouflage coating for leafhoppers"

**This PDF file includes:**

Figures S1 to S8

Tables S1 and S2

17 **Supplementary Figures 1-6**

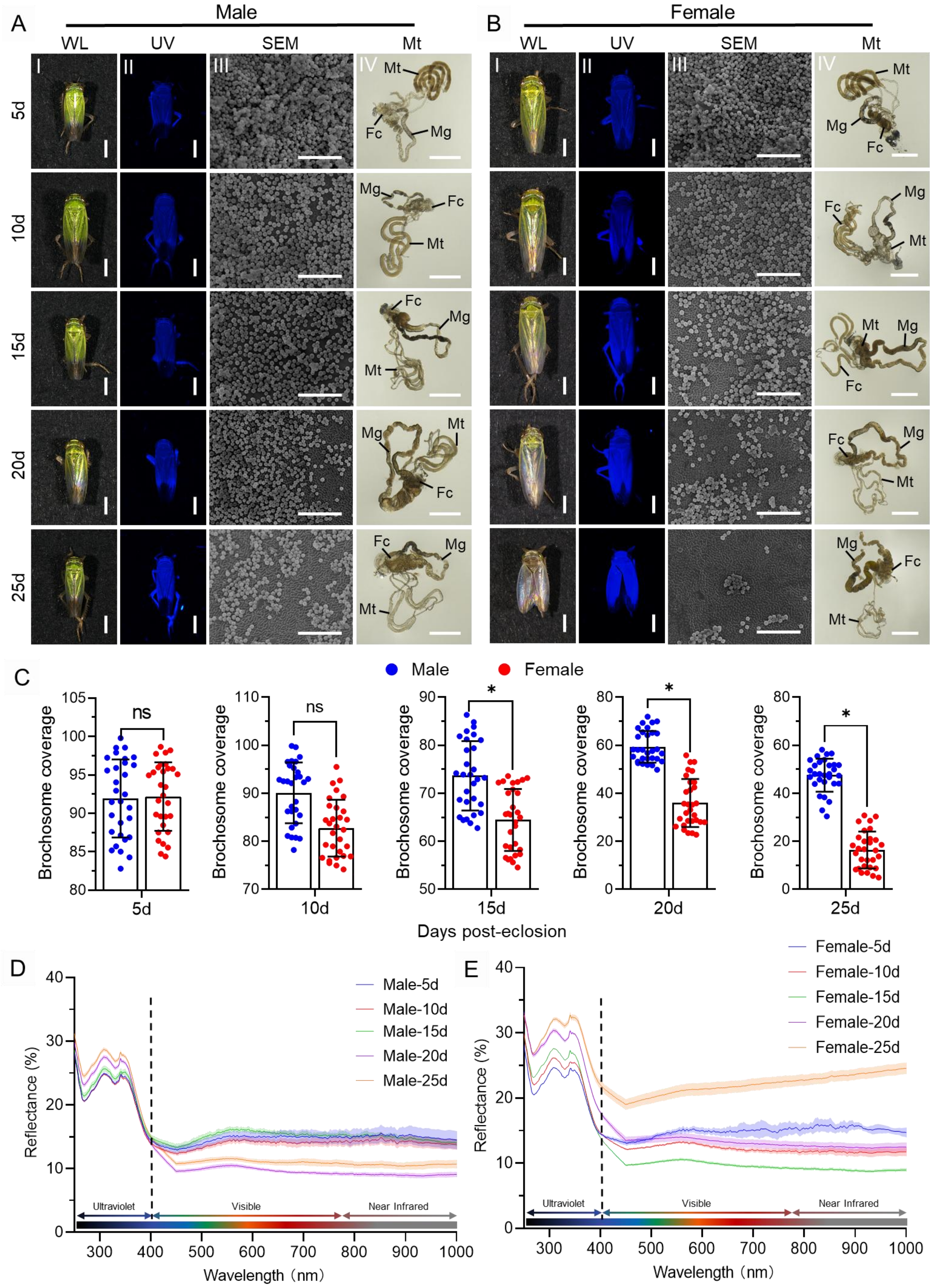


18

**Figure S1: Influence of brochosome coating on optical properties of *N. cincticeps* cuticle surface.** (A, B) Gradual reduction of brochosome coverage on the cuticle surface of *N. cincticeps* post-eclosion. Images were captured depicting male (A) and female (B) *N. cincticeps* over the course of 5 to 25 days post-eclosion under both white and ultraviolet light. SEM was employed to scrutinize the distribution of microstructures on the cuticle surface during this period. Additionally, dissections were carried out to observe and document morphological changes in the Malpighian tubules of leafhoppers *N. cincticeps* over the same timeframe. All images are representative of at least three replicates. Bar, 1 mm in I, II, and IV; 5 μm in III. (C) The distribution area ratio of brochosomes on the cuticle surface of *N. cincticeps* was statistically analyzed at 5, 10, 15, 20, and 25 days post-eclosion. Each data point represents the outcome of an individual independent experiment. The presented data are expressed as mean ± SD values. Statistical significance is denoted as *$P < 0.05$ and ns no significance, determined by unpaired t test. (D-E) Reflectance spectra of male (D) and female (E) forewing of *N. cincticeps* from 5 to 25 days post-eclosion. All images are representative of at least three replicates.

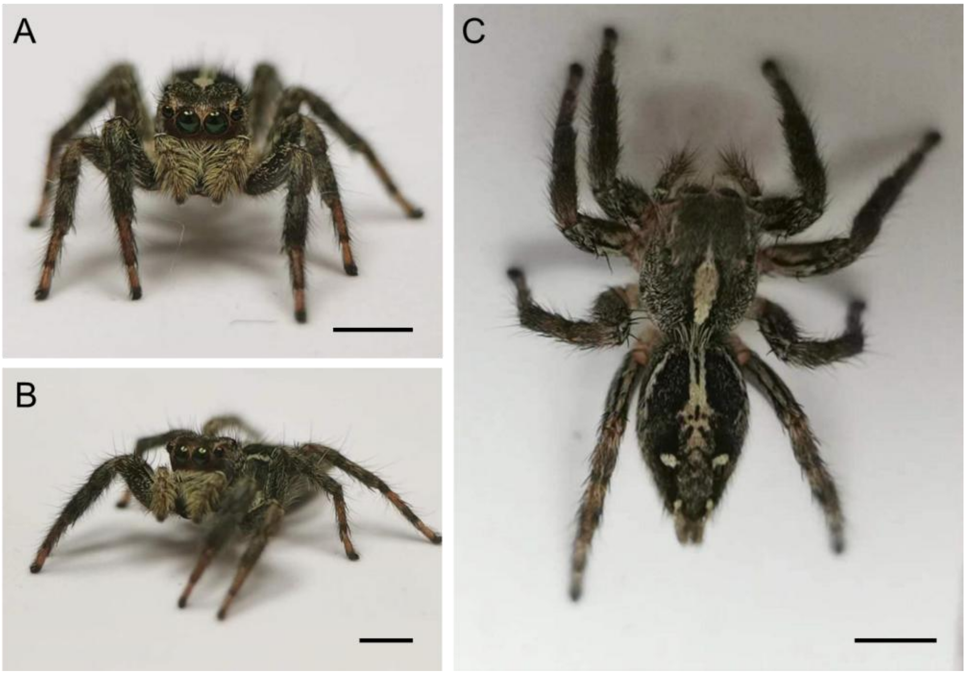


**Figure S2: Frontal view (A), top view (B) and side view (C) of jumping spider *P. paykulli*.**

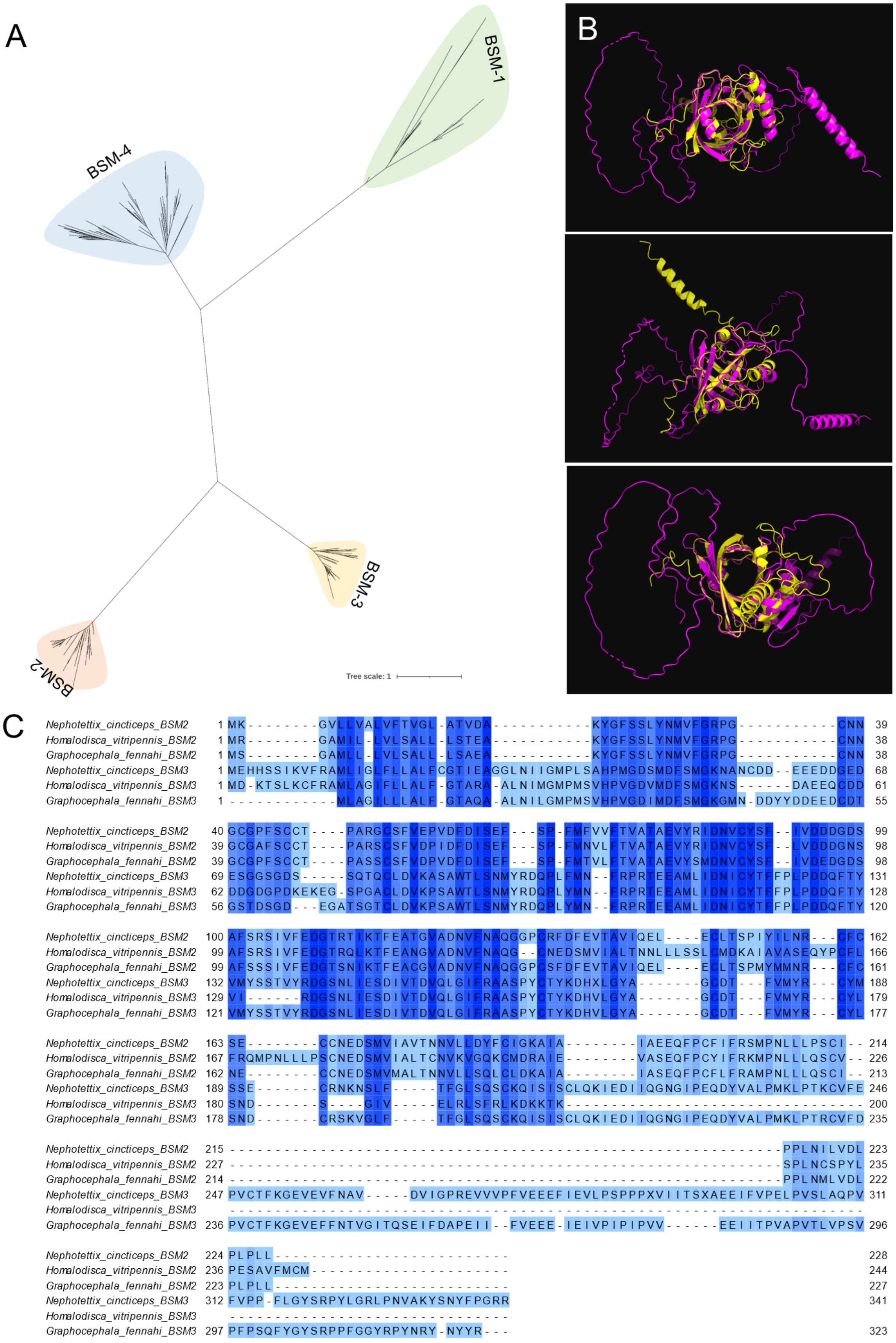
A
B
C
BSM-1
BSM-2
BSM-3
BSM-4
Tree scale: 1
Nephotettix_cincticeps_BSM2 1 MK--------GVLLVALVFTVGL-ATVDA-----------KYGFSSLYNMVFGRPG-----------CNN 39
Homalodisca_vitripennis_BSM2 1 MR--------GAMIL-LVLSALL-LSTEA-----------KYGFSSLYNMVFGRPG-----------CNN 38
Graphocephala_fennahi_BSM2 1 MS--------GAMLL-LVLSALL-LSAEA-----------KYGFSSLYNMVFGRPG-----------CNN 38
Nephotettix_cincticeps_BSM3 1 MEHHSSIKVFRAMLIGLFLLALFCGTIEAGGLNIIGMPLSAHPMGDSMDFSMGKNANCDD--EEEDDGED 68
Homalodisca_vitripennis_BSM3 1 MD-KTSLKCFRAMLAGIFLLALF-GTARA-ALNIMGMPMSVHPVGDVMDFSMGKNS------DAEEQCDD 61
Graphocephala_fennahi_BSM3 1 ------------MLAGILLLALF-GTAQA-ALNILGMPMSVHPVGDIMDFSMGKGMN-DDYYDDEEDCDT 55
Nephotettix_cincticeps_BSM2 40 GCGPFSCCT----PARGCSFVEPVDFDISEF---SP-FMFVVFTVATAEVYRIDNVCYSF--IVDDDGDS 99
Homalodisca_vitripennis_BSM2 39 GCGAFSCCT----PARSCSFVDPIDFDISEF---SP-FMNVLFTVATAEVYRIDNVCYSF--LVDDDGNS 98
Graphocephala_fennahi_BSM2 39 GCGPFSCCT----PASSCSFVDPVDFDISEF---SP-FMTVLFTVATAEVYSMDNVCYSF--IVDEDGDS 98
Nephotettix_cincticeps_BSM3 69 ESGGSGDS-----SQTQCLDVKASAWTLSNMYRDQPLFMN--FRPRTEEAMLIDNICYTFFPLPDDQFTY 131
Homalodisca_vitripennis_BSM3 62 DDGDGPDKEKEG-SPGACLDVKPSAWTLSNMYRDQPLYMN--FRPRTEEAMLIDNICYTFFPLPDDQFTY 128
Graphocephala_fennahi_BSM3 56 GSTDSGD---EGATSGTCLDVKPSAWTLSNMYRDQPLYMN--FRPRTEEAMLIDNICYTFFPLPDDQFTY 120
Nephotettix_cincticeps_BSM2 100 AFSRSIVFEDGTRTIKTFEATGVADNVFNAQGGPCRFDFEVTAVIQEL----ECLTSPIYILNR---CFC 162
Homalodisca_vitripennis_BSM2 99 AFSRSIVFEDGTRQLKTFEANGVADNVFNAQG--CNEDSMVIALTNNLLLSSLCMDKAIAVASEQYPCFL 166
Graphocephala_fennahi_BSM2 99 AFSSSIVFEDGTSNIKTFEACGVADNVFNAQGGPCSFDFEVTAVIQEL----ECLTSPMYMMNR---CFC 161
Nephotettix_cincticeps_BSM3 132 VMYSSTVYRDGSNLIESDIVTDVQLGIFRAASPYCTYKDHXLGYA-------GCDT---FVMYR---CYM 188
Homalodisca_vitripennis_BSM3 129 VI------RDGSNLIESDIVTDVQLGIFRAASPYCTYKDHVLGYA-------GCDT---FVMYR---CYL 179
Graphocephala_fennahi_BSM3 121 VMYSSTVYRDGSNLIESDIVTDVQLGIFRAASPYCTYKDHVLGYA-------GCDT---FVMYR---CYL 177
Nephotettix_cincticeps_BSM2 163 SE--------CCNEDSMVIAVTNNVLLDYFCIGKAIA--------IAEEQFPCFIFRSMPNLLLPSCI-- 214
Homalodisca_vitripennis_BSM2 167 FRQMPNLLLPSCNEDSMVIALTCNVKVGQKCMDRAIE--------VASEQFPCYIFRKMPNLLLQSCV-- 226
Graphocephala_fennahi_BSM2 162 NE--------CCNEDSMVMALTNNVLLSQLCLDKAIA--------IASEQFPCFLFRAMPNLLLQSCI-- 213
Nephotettix_cincticeps_BSM3 189 SSE-------CRNKNSLF-----TFGLSQSCKQISISCLQKIEDIIQGNGIPEQDYVALPMKLPTKCVFE 246
Homalodisca_vitripennis_BSM3 180 SND-------S---GIV-----ELRLSFRLKDKKTK----------------------------------- 200
Graphocephala_fennahi_BSM3 178 SND-------CRSKVGLF-----TFGLSQSCKQISISCLQKIEDIIQGNGIPEQDYVALPMKLPTRCVFD 235
Nephotettix_cincticeps_BSM2 215 -------------------------------------------------------------PPLNILVDL 223
Homalodisca_vitripennis_BSM2 227 -------------------------------------------------------------SPLNCSPYL 235
Graphocephala_fennahi_BSM2 214 -------------------------------------------------------------PPLNMLVDL 222
Nephotettix_cincticeps_BSM3 247 PVCTFKGEVEVFNAV----DVIGPREVVVPFVEEFIEVLPSPPPXVIITSXAEEIFVPELPVSLAQPV 311
Homalodisca_vitripennis_BSM3 ----------------------------------------------------------------------
Graphocephala_fennahi_BSM3 236 PVCTFKGEVEFFNTVGITQSEIFDAPEII--FVEEE-IEIVPIPIPVV------EEIITPVAPVTLVPSV 296
Nephotettix_cincticeps_BSM2 224 PLPLL-------------------------- 228
Homalodisca_vitripennis_BSM2 236 PESAVFMCM---------------------- 244
Graphocephala_fennahi_BSM2 223 PLPLL-------------------------- 227
Nephotettix_cincticeps_BSM3 312 FVPP-FLGYSRPYLGRLPNVAKYSNYFPGRR 341
Homalodisca_vitripennis_BSM3 -------------------------------
Graphocephala_fennahi_BSM3 297 PFPSQFYGYSRPPFGGYRPYNRY-NYYR--- 323

37

**Figure S3: Homology analysis of BSM-encoding genes.** **(A)** Maximum likelihood phylogenetic tree based on the four BSM-encoding genes was constructed with a bootstrap of 1000.(**B)** Comparison of protein tertiary structures of BSM2 and BSM3. Yellow color indicates BSM2, pink color indicates BSM3. (**C)** Alignment of amino acid sequences of BSM2 and BSM3 identified in *N. cincticeps* and laccases of other two Cicadellidae species.

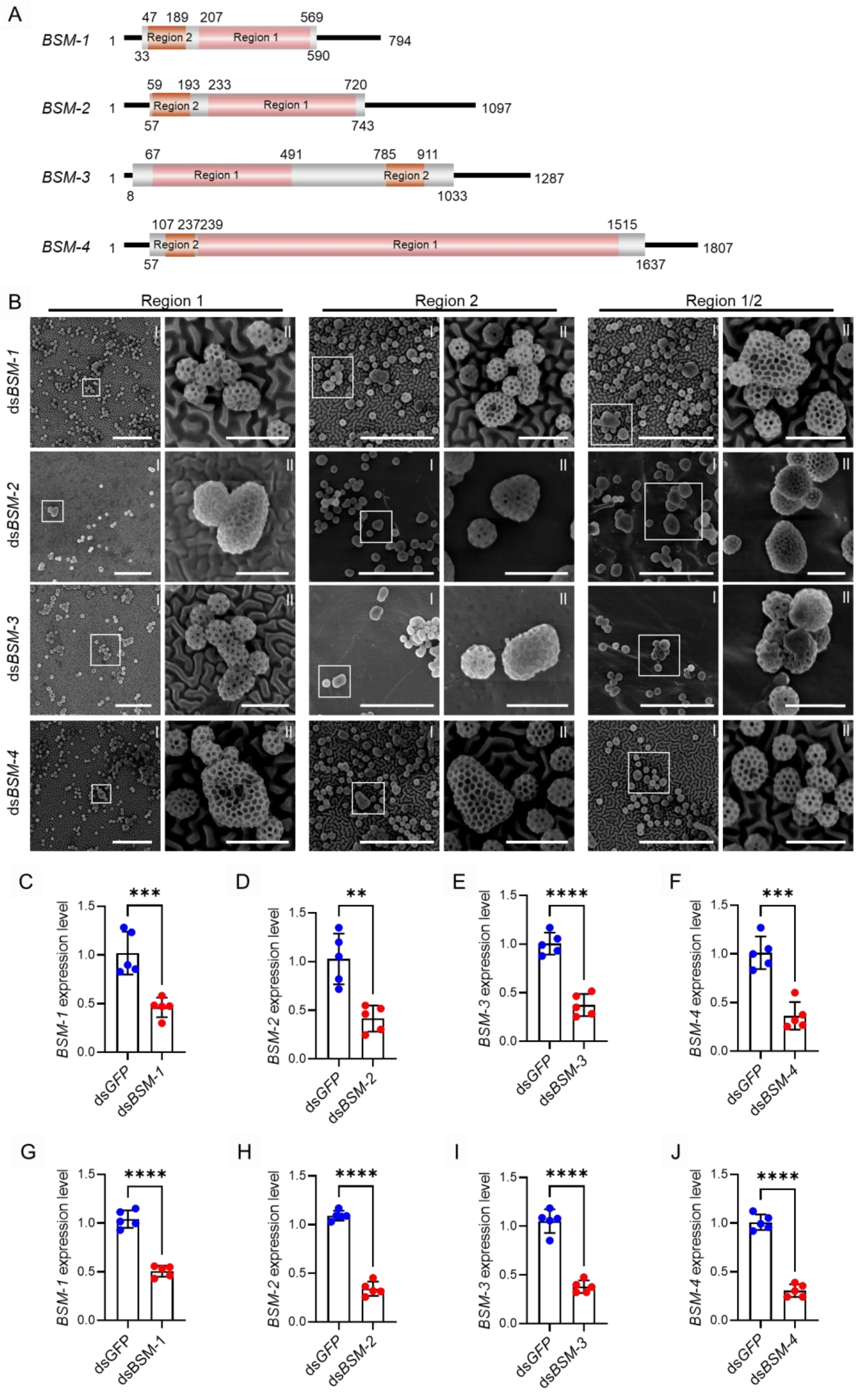

A
BSM-1
BSM-2
BSM-3
BSM-4
Region 1
Region 2
B
Region 1
Region 2
Region 1/2
dsBSM-1
dsBSM-2
dsBSM-3
dsBSM-4
C
D
E
F
G
H
I
J
BSM-1 expression level
BSM-2 expression level
BSM-3 expression level
BSM-4 expression level
dsGFP
dsBSM-1
dsBSM-2
dsBSM-3
dsBSM-4


44

**Figure S4: BSM1-4 are brochosomal structural proteins.** (A) Distribution of dsRNAs targeting two non-overlapping regions in four BSM-encoding genes. (B) Brochosome morphology on N. cincticeps forewings at 7 days post-microinjection of dsRNA targeting two non-overlapping regions of each BSM-encoding gene. II are partial enlarged images for I. Scale bar in I, 5 μm; II, 1 μm. (C-J) Transcription levels of BSM-1 (C, Region1; G, Region2), BSM-2 (D, Region1; H, Region2), BSM-3 (E, Region1; I, Region2), and BSM-4 (F, Region1; J, Region2) at 7 days post-microinjection of dsRNA targeting two non-overlapping regions of each BSM-encoding gene.

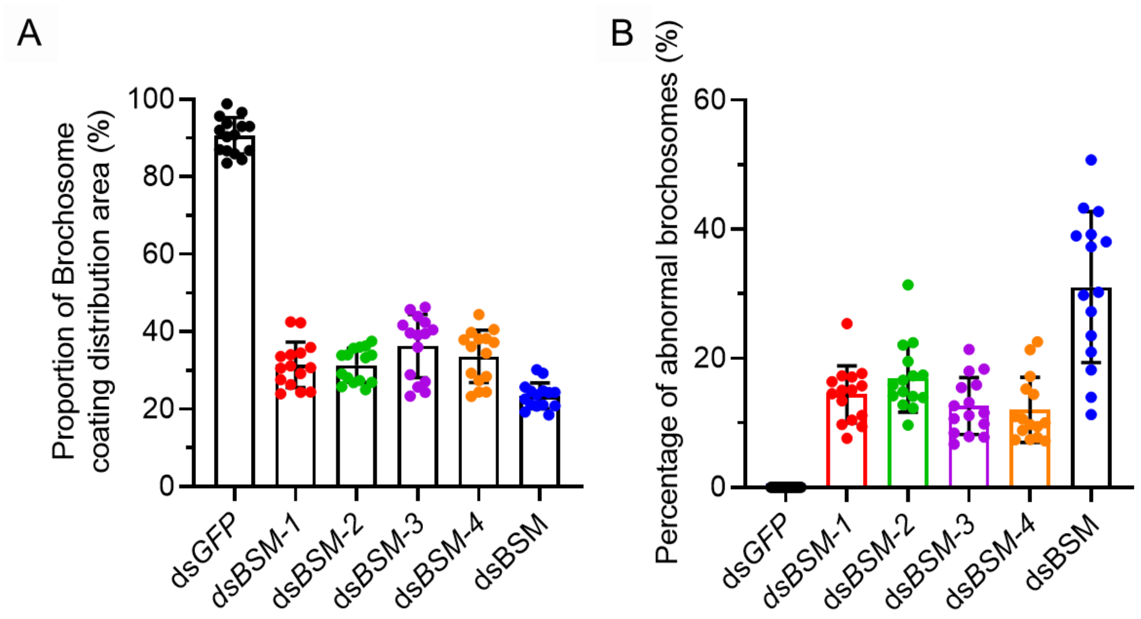


**Figure S5: Inhibition of BSM gene expression reduces brochosome distribution on *N. cincticeps* cuticle surface (A) and alters their morphology (B).**

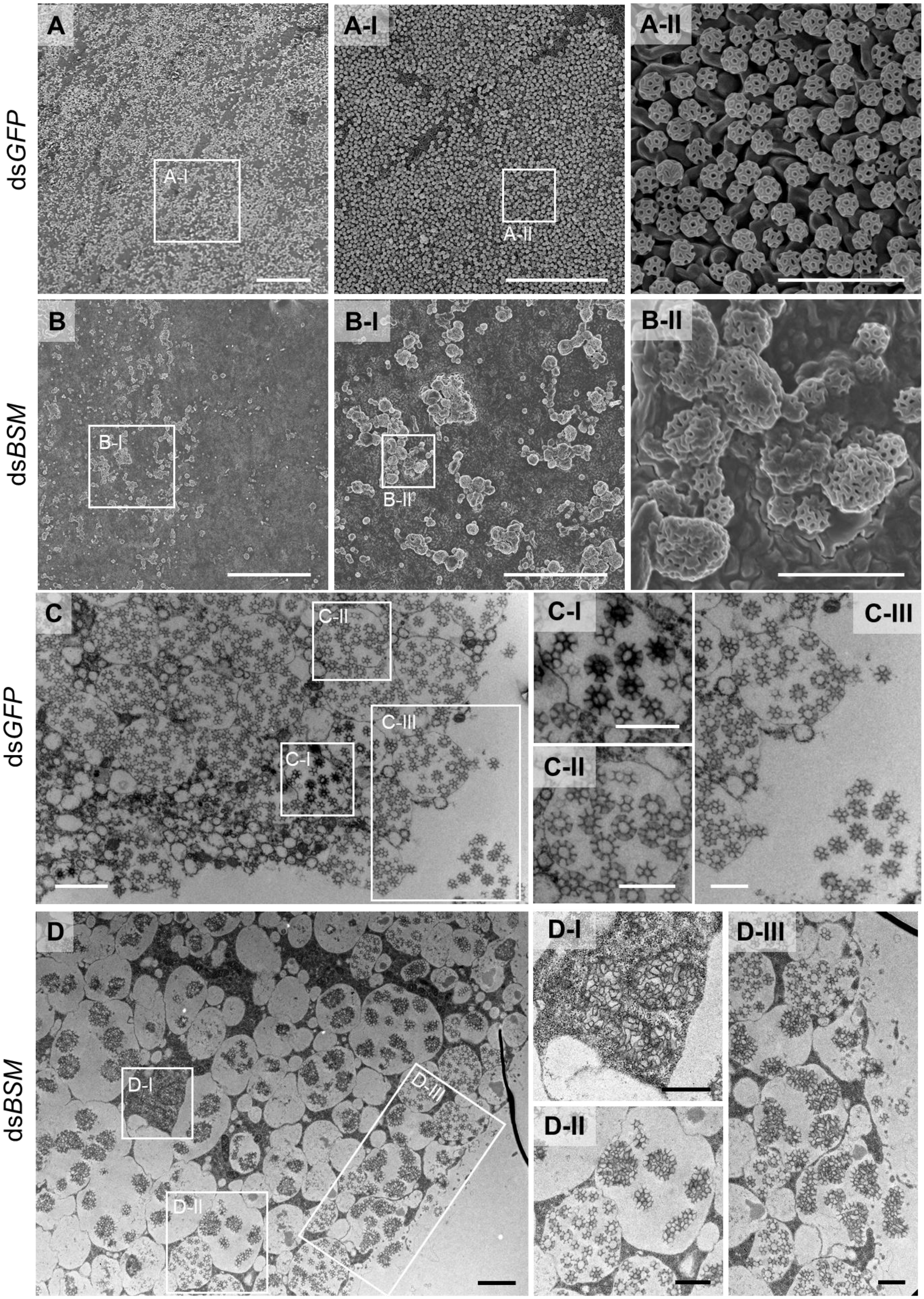

dsGFP
dsBSM
dsGFP
dsBSM
A
A-I
A-II
B
B-I
B-II
C
C-I
C-II
C-III
D
D-I
D-II
D-III

**Figure S6: Differences in the synthesis, morphology, and distribution of brochosomes following RNAi treatment.** (A, B) Morphology of the brochosome on the forewings of leafhoppers after ds*GFP* (A) and ds*BSM* (B) treatment. Images in A-I and A-II represent successive magnifications of the boxed region in A and A-I, respectively. Similarly, images in B-I and B-II depict successive magnifications of the boxed region in B and B-I, respectively. Scale bar in A and B, 25 μm; A-I and B-I, 10 μm; A-II and B-II, 1 μm. (C and D) Synthesis and morphology of the brochosome in the distal segment epithelial cells of Malpighian tubules after ds*GFP* (C) and ds*BSM* (D) treatment. C-I, C-II, C-III, D-I, D-II, and D-III are enlargements of the boxed regions in C and D, respectively.Scale bars in C-I, C-II, C-III, D-I, D-II, and D-III, 1 μm; C and D, 2 μm.

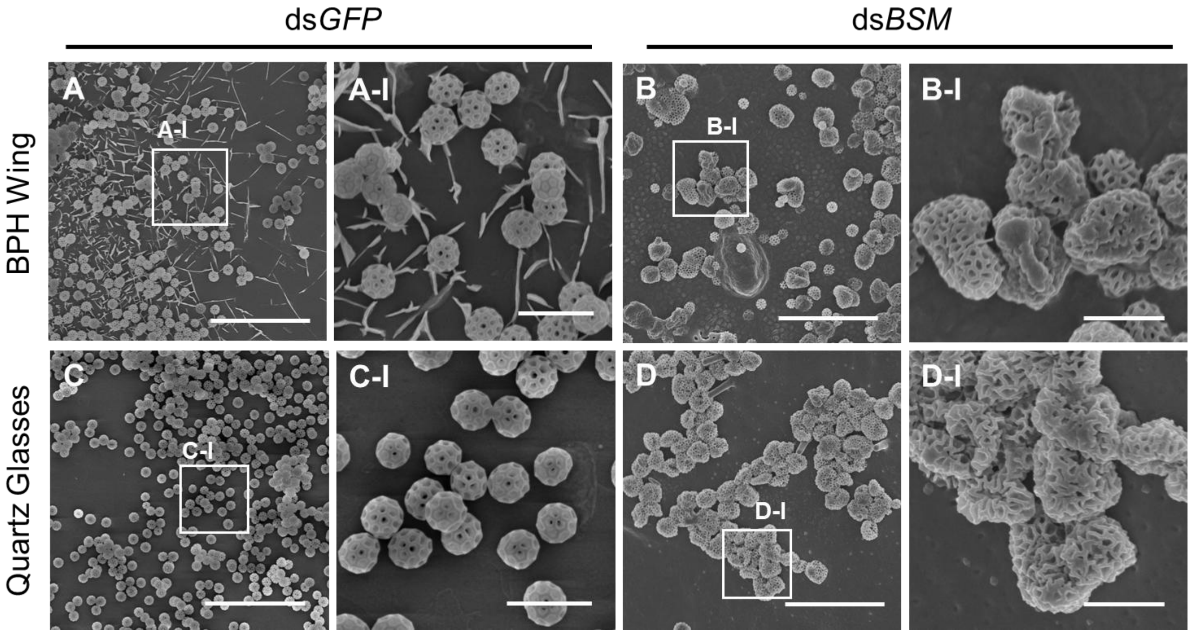


**Figure S7: The distribution of brochosomes, collected after ds*GFP* and ds*BSM* treatments, applied to both the wings of BPH (A, B) and quartz glass (C, D).** A-I, B-I, C-I, D-I are enlargements of the boxed regions in A, B, C, and D, respectively. Scale bars in A-D, 5 μm; A-I, B-I, C-I, and D-I, 1 μm.

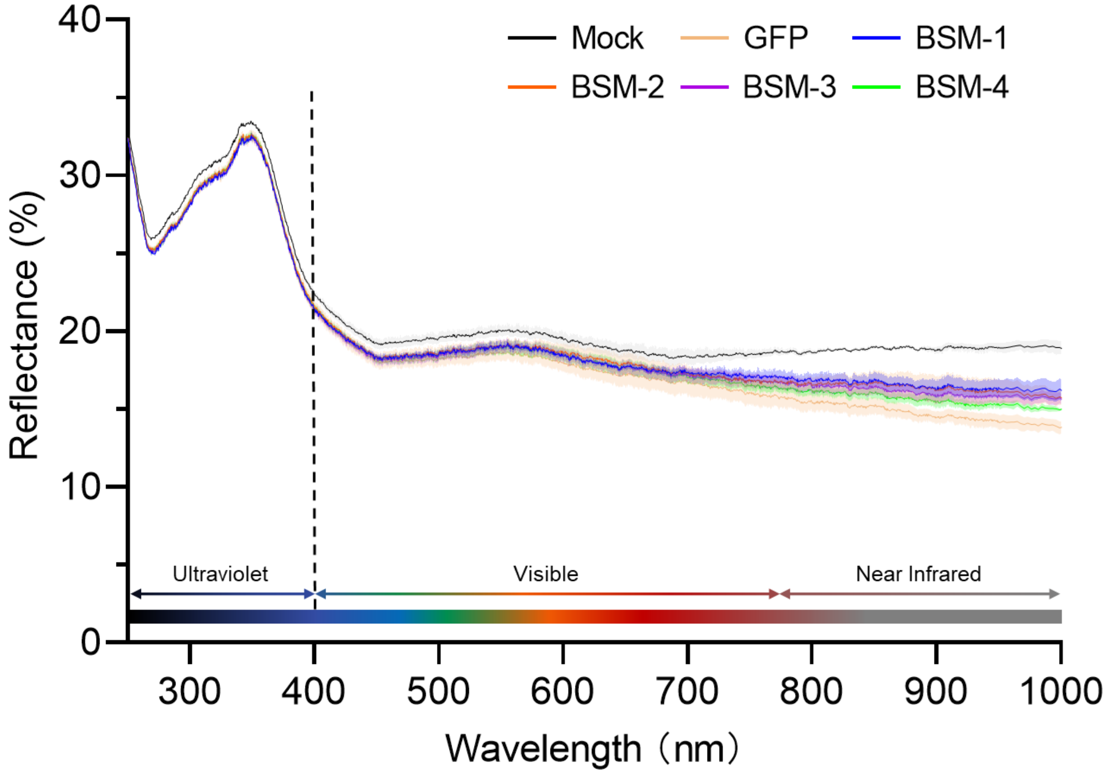


**Figure S8: The purified BSM protein lacks anti-reflective properties. Quartz slides treated with purified GST protein were used as the negative control, and those treated with PBS (phosphate-buffered saline) alone served as the mock.**

**Table S1. List of oligonucleotide primers used in this study**

| Oligonucleotide | Assay | Sequence (5'-3') |
|---|---|---|
| CO1_F | PCR | TTGATTTTTTGGTCATCCAGAAGT |
| CO1_R | PCR | TCCAATGCACTAATCTGCCATATTA |
| qPCR_BSM1_F | qPCR | CTCCTTGACCGCTTCATC |
| qPCR_BSM1_R | qPCR | CGCCATCCTGTTCTTCAT |
| qPCR_BSM2_F | qPCR | CAGTTCCCTTGCTTCATTT |
| qPCR_BSM2_R | qPCR | CCACCAGGATGTTCAGAG |
| qPCR_BSM3_F | qPCR | GATGACCAGTTCACCTAC |
| qPCR_BSM3_R | qPCR | GTCCGTAACAATGTCTGA |
| qPCR_BSM4_F | qPCR | ACTGGTGGACAACATGAA |
| qPCR_BSM4_R | qPCR | AAGTAGAGATGCCGTTGAT |
| qPCR_EF1_F | qPCR | CAGTGAGAGCCGTTTTGAG |
| qPCR_EF1_R | qPCR | AGGGCATCTTGTCAGAGGGC |
| T7_BSM1_ region1_F | RNAi | GGATCCTAATACGACTCACTATAGGCTCCTT<br>GACCGCTTCATC |
| T7_BSM1_ region1_R | RNAi | GGATCCTAATACGACTCACTATAGGGTCAC<br>TATCTCCAAGGTT |
| T7_BSM2_ region1_F | RNAi | GGATCCTAATACGACTCACTATAGGCGACT<br>TTGACATATCTGAG |
| T7_BSM2_ region1_R | RNAi | GGATCCTAATACGACTCACTATAGGCCACC<br>AGGATGTTCAGAG |
| T7_BSM3_ region1_F | RNAi | GGATCCTAATACGACTCACTATAGGATGCT<br>TATCGGCCTCTTTCTTTTG |
| T7_BSM3_ region1_R | RNAi | GGATCCTAATACGACTCACTATAGGCTGCT<br>CTGAAGATTCCAA |
| T7_BSM4_ region1_F | RNAi | GGATCCTAATACGACTCACTATAGGGTCAC<br>TGGTGGACAACAT |
| T7_BSM4_ region1_R | RNAi | GGATCCTAATACGACTCACTATAGGTTGTA<br>GAAGGCGTGTAAGT |
| T7_BSM1_region2_F | RNAi | GGATCCTAATACGACTCACTATAGGTGTTCT<br>CGCTGTTTTCACCG |
| T7_BSM1_region2_R | RNAi | GGATCCTAATACGACTCACTATAGGTTAGA<br>CGATCCAACCATGCG |
| T7_BSM2_region2_F | RNAi | GGATCCTAATACGACTCACTATAGGGAAAG<br>GGGTGTTGTTGGTGG |

| | | |
|---|---|---|
| T7_BSM2_region2_R | RNAi | GGATCCTAATACGACTCACTATAGGCAGCTGAAAGGACCGCAG |
| T7_BSM3_region2_F | RNAi | GGATCCTAATACGACTCACTATAGGGCTGTGGATGTCATTGGTCC |
| T7_BSM3_region2_R | RNAi | GGATCCTAATACGACTCACTATAGGCAGGGACGAAGATCTCCTCG |
| T7_BSM4_region2_F | RNAi | GGATCCTAATACGACTCACTATAGGTCGAGCGGAGATAAATGTGAA |
| T7_BSM4_region2_R | RNAi | GGATCCTAATACGACTCACTATAGGCTGTCTCCGATGCCTTCTGT |
| T7_GFP_F | RNAi | GGATCCTAATACGACTCACTATAGGCACAAGTTCAGCGTGTCCG |
| T7_GFP_R | RNAi | GGATCCTAATACGACTCACTATAGGGTTCACCTTGATGCCGTTC |
| RACE_BSM1_F1 | RACE | GATTACGCCAAGCTTCCTTGACCGCTTCATCGCCCTCATACCC |
| RACE_BSM1_R1 | RACE | GATTACGCCAAGCTTGCCTTCGTCACTGCCGCTGTCACCTTC |
| RACE_BSM1_F2 | RACE | GATTACGCCAAGCTTCTGAGTGCGTATATCAAGGGCGTAGT |
| RACE_BSM1_R2 | RACE | GATTACGCCAAGCTTCTGTCACCTTCCTCTTCGGATGTAGTC |
| RACE_BSM2_F1 | RACE | GATTACGCCAAGCTTTGGTGGCGTTGGTGTTCACTGTCGGCTT |
| RACE_BSM2_R1 | RACE | GATTACGCCAAGCTTGAGCATCCTCTAGCGGGCGTGCAGCAG |
| RACE_BSM2_F2 | RACE | GATTACGCCAAGCTTGCCAAGTACGGATTCTCCTCTCTGT |
| RACE_BSM2_R2 | RACE | GATTACGCCAAGCTTCTGAAAGGACCGCAGCCGTTGTTG |
| RACE_BSM3_F1 | RACE | GATTACGCCAAGCTTGCTGGCGGCTTGAACATCATCGGCATG |
| RACE_BSM3_R1 | RACE | GATTACGCCAAGCTTCGTAGGTGAACTGGTCATCCGGCAGAGG |
| RACE_BSM3_F2 | RACE | GATTACGCCAAGCTTACGAAGAGGAAGACGATGGCGAGGAT |

| | | |
|---|---|---|
| RACE_BSM3_R2 | RACE | GATTACGCCAAGCTTCGGCAGAGGGAAGAAGGTATAGCAGATG |
| RACE_BSM4_F1 | RACE | GATTACGCCAAGCTTCATCGACACCTGTGCCCGCACCTACCC |
| RACE_BSM4_R1 | RACE | GATTACGCCAAGCTTGACACGACTCCATTCGCCGCCGTCTGG |
| RACE_BSM4_F2 | RACE | GATTACGCCAAGCTTGGAATCGTTTGAGGACATGCGTTG |
| RACE_BSM4_R2 | RACE | GATTACGCCAAGCTTCTCCCCACAGTGCTTGCTACCAC |

**Table S2. Information on hemipteran insects used in the search for homologs of BSM**

| Suborder | Family | Organism | BioProject |
|---|---|---|---|
| Coleorrhyncha | Peloridiidae | Hackeriella veitchi | PRJNA357411 |
| Sternorrhyncha | Aphrophoridae | Philaenus spumarius | PRJNA272277 |
| | Aphrophoridae | Aphrophora alni | PRJNA272162 |
| | Cercopidae | Prosapia bicincta | PRJNA272284 |
| | | Cercopis vulnerata | PRJNA183205 |
| | Clastopteridae | Clastoptera obtusa | PRJNA295707 |
| | | Clastoptera arizonana | PRJNA303152 |
| | Epipygidae | Epipyga sp. | PRJNA295739 |
| | Machaerotidae | Pectinariophyes stalii | PRJNA295734 |
| Auchenorrhyncha | Cicadidae | Maoricicada tenuis | PRJNA295717 |
| | | Kikihia scutellaris | PRJNA295715 |
| | | Guineapsaltria flava | PRJNA295700 |
| | | Tamasa doddi | PRJNA295701 |
| | | Megatibicen dorsatus | PRJNA272295 |
| | | Burbunga queenslandica | PRJNA295699 |
| | | Tettigades auropilosa | PRJNA295726 |
| | | Okanagana villosa | PRJNA183205 |
| | | Chilecicada sp. | PRJNA295722 |
| | Peloridiidae | Xenophysella greensladeae | PRJNA183205 |
| | | Xenophyes metoponcus | PRJNA272209 |
| | | Peloridium pomponorum | PRJNA272276 |
| | Tettigarctidae | Tettigarcta crinita | PRJNA295711 |
| | Acanaloniidae | Acanalonia conica | PRJNA272210 |
| | Achilidae | Catonia nava | PRJNA295706 |
| | Caliscelidae | Caliscelis bonellii | PRJNA272168 |
| | | Bruchomorpha oculata | PRJNA272222 |
| | Cixiidae | Tachycixius pilosus | PRJNA272206 |
| | | Melanoliarus placitus | PRJNA272269 |
| | Delphacidae | Idiosystatus acutiusculus | PRJNA272251 |
| | | Nilaparvata lugens | PRJNA183205 |
| | Derbidae | Omolicna uhleri | PRJNA295709 |
| | Dictyopharidae | Yucanda albida | PRJNA357446 |
| | | Phylloscelis atra | PRJNA272279 |
| | | Dictyophara europaea | PRJNA272176 |

| | | |
|---|---|---|
| | Chondrodire chilensis | PRJNA272294 |
| Eurybrachidae | Platybrachys sp. | PRJNA295736 |
| Flatidae | Ormenoides venusta | PRJNA272272 |
| | Metcalfa pruinosa | PRJNA272198 |
| | Jamella australiae | PRJNA295743 |
| | Geisha distinctissima | PRJNA352589 |
| Fulgoridae | Lycorma delicatula | PRJNA357415 |
| | Pyrops candelaria | PRJNA352589 |
| | Cyrpoptus belfragei | PRJNA272237 |
| Issidae | Gergithus sp. | PRJNA352589 |
| | Thionia simplex | PRJNA295721 |
| Nogodinidae | Lipocallia australensis | PRJNA295732 |
| | Bladina sp. | PRJNA357426 |
| Ricaniidae | Scolypopa sp. | PRJNA295737 |
| | Ricania speculum | PRJNA357442 |
| Tettigometridae | Tettigometra bipunctata | PRJNA357421 |
| Tropiduchidae | Ladella sp. | PRJNA272255 |
| Aetalionidae | Lophyraspis sp. | PRJNA357414 |
| | Aetalion reticulatum | PRJNA357422 |
| Cicadellidae | Agallia constricta | PRJNA272213 |
| | Aphrodes bicincta | PRJNA357425 |
| | Vidanoana flavomaculata | PRJNA272302 |
| | Homalodisca vitripennis | PRJNA415461; PRJNA342859 |
| | Homalodisca liturata | PRJNA303151 |
| | Graphocephala fennahi | PRJNA272183 |
| | Graphocephala coccinea | PRJNA341855 |
| | Graphocephala atropunctata | PRJNA299492 |
| | Cuerna arida | PRJNA303150 |
| | Cicadella viridis | PRJNA629998 |
| | Tinobregmus viridescens | PRJNA295727 |
| | Scaphoideus titanus | PRJNA765507 |
| | Recilia dorsalis | PRJNA629998 |
| | Psammotettix striatus | PRJNA743281 |
| | Penthimia sp. | PRJNA357438 |
| | Nephotettix virescens | PRJNA671146 |
| | Neoaliturus tenellus | PRJNA318850 |

| | | |
|---|---|---|
| | Macrosteles quadrilineatus | PRJNA562189; PRJNA438515;PRJNA318848 |
| | Haldorus sp. | PRJNA357443 |
| | Graminella nigrifrons | PRJNA167489 |
| | Exitianus exitiosus | PRJNA562189 |
| | Exitianus capicola | PRJNA807480 |
| | Euscelidius variegatus | PRJNA393620 |
| | Euacanthella palustris | PRJNA295730 |
| | Dalbulus maidis | PRJNA505618; PRJNA318849; PRJNA631706; PRJNA272239 |
| | Balclutha rubrostriata | PRJNA562189 |
| | Balclutha neglecta | PRJNA562189 |
| | Agudus sp. AgspCi3 | PRJNA357423 |
| | Macropsis decisa | PRJNA295724 |
| | Idiocerus rotundens | PRJNA295723 |
| | Ponana quadralaba | PRJNA272282 |
| | Penestragania robusta | PRJNA357439 |
| | Tituria crinita | PRJNA357436 |
| | Stenocotis depressa | PRJNA295738 |
| | Hespenedra chilensis | PRJNA272247 |
| | Neocoelidia tumidifrons | PRJNA295725 |
| | Nionia palmeri | PRJNA295720 |
| | Zyginidia pullula | PRJNA171390 |
| | Watara sudra | PRJNA718899 |
| | Matsumurasca onukii | PRJNA347531 |
| | Empoasca fabae | PRJNA272241 |
| | Amrasca biguttula biguttula | PRJNA555157 |
| | Ulopa reticulata | PRJNA272207 |
| | Xestocephalus desertorum | PRJNA295729 |
| Melizoderidae | Llanquihuea pilosa | PRJNA272258 |
| | Umbonia crassicornis | PRJNA295728 |
| | Tolania sp. | PRJNA357445 |
| | Stictocephala bisonia | PRJNA272293 |
| | Procyrta sp. | PRJNA357441 |
| | Notocera sp. | PRJNA357437 |
| | Nessorhinus gibberulus | PRJNA272268 |
| | Microcentrus caryae | PRJNA295719 |

| | | |
|---|---|---|
| | Membracis tectigera | PRJNA357433 |
| | Lycoderes burmeisteri | PRJNA357432 |
| | Holdgatiella chepuensis | PRJNA272249 |
| | Heteronotus sp. | PRJNA295741 |
| | Entylia carinata | PRJNA415461 |
| | Enchenopa latipes | PRJNA272226 |
| | Cyphonia clavata | PRJNA357428 |
| | Chelyoidea sp. | PRJNA357427 |
| | Centrotus cornutus | PRJNA272169 |
| | Amastris sp. | PRJNA357424 |
| | Mapuchea sp. | PRJNA272263 |
